# Transcriptomic metaanalyses of autistic brains reveals shared gene expression and biological pathway abnormalities with cancer

**DOI:** 10.1101/437905

**Authors:** Jaume Forés-Martos, Ferrán Catalá-López, Jon Sánchez-Valle, Kristina Ibáñez, Héctor Tejero, Helena Palma-Gudiel, Joan Climent, Vera Pancaldi, Lourdes Fañanás, Celso Arango, Mara Parellada, Anaïs Baudot, Daniel Vogt, John L. Rubenstein, Alfonso Valencia, Rafael Tabarés-Seisdedos

## Abstract

Epidemiological and clinical evidence points to cancer as a comorbidity in people with autism spectrum disorders (ASD). A significant overlap of genes and biological processes between both diseases has also been reported. Here, for the first time, we compared the gene expression profiles of ASD frontal cortex tissues and 22 cancer types obtained by differential expression meta-analysis. Four cancer types (brain, thyroid, kidney, and pancreatic cancers) presented a significant overlap in gene expression deregulations in the same direction as ASD whereas two cancer types (lung and prostate cancers) showed differential expression profiles significantly deregulated in the opposite direction from ASD. Functional enrichment and LINCS L1000 based drug set enrichment analyses revealed the implication of several biological processes and pathways that were affected jointly in both diseases, including impairments of the immune system, and impairments in oxidative phosphorylation and ATP synthesis among others. Our data also suggest that brain and kidney cancer have patterns of transcriptomic dysregulation in the PI3K/AKT/MTOR axis that are similar to those found in ASD. These shared transcriptomic alterations could help explain epidemiological observations suggesting direct and inverse comorbid associations between ASD and particular cancer types.

## Introduction

As Jane Austin once wrote, *It is a truth universally acknowledged that*(1) autism spectrum disorder (ASD) is a chronic childhood-onset neurodevelopmental condition characterized by persistent deficits in social communication and social interactions, as well as, by restricted, repetitive patterns of behavior, interests, or activities(2, 3). However, other serious clinical aspects of ASD are less well known. For instance, an increase in premature mortality has been recently reported(4–7). ASD is among the top ten causes of disability worldwide in children between 5 and 9 years old(8), these findings could be partially explained by the link between ASD and other lifetime health problems, including epilepsy, diabetes, cardiovascular and gastrointestinal diseases, cancer, depression, and suicide(8–12). A better understanding of these lifetime co-occurring conditions is important for people with ASD, their families and caregivers, clinicians and other healthcare professionals, scientists, and policy makers(13, 14). Recognizing this multimorbidity scenario, we focus our attention on the relationships between ASD and cancer for two reasons. First, evidence pointing towards different cancer rates in patients with central nervous system disorders has started to been gathered(15). Although several studies have failed to find specific associations between ASD and cancer(16–18), others, including a large population cohort study in Taiwan(11) suggested a higher-than-expected occurrence of overall cancer in ASD patients. These authors found a standardized incidence ratio of 1.94 (95% CI 1.18-2.99), with further increased incidence for brain and genitourinary cancers. Similarly, a large population-based case-control study in Sweden noted a significant increase in cancer mortality for all cancers combined (OR = 1.80, 95% CI 1.46-2.23) among individuals with ASD as compared with the general population(6). In addition, mothers of children with ASD have been shown to be approximately 50% more likely to die from cancer than those of non-autistic offspring(17). Conversely, two studies found a lower-than-expected risk of neoplasm in ASD patients, a situation that could be described as “inverse cancer comorbidity”(12, 19). Second, given the prevalence and social impact of both diseases, further characterization of the genetic, molecular and cellular factors involved in ASD and cancer, which represent their underlying mechanisms and are used in their identification, are important and incompletely resolved issues. Recent genome-wide exome-sequencing studies of *de novo* variants and recurrent copy number variations (CNVs) in ASD and cancer have revealed extensive overlap in risk genes for autism and cancer(20–24). Moreover, several studies have found a striking implication of the classically cancer related PTEN pathway in ASD(22, 25–27). These findings provide persuasive evidence of a molecular link between ASD and cancer, possibly opening the door to new treatments for both conditions. For example, chemotherapeutic agents that inhibit PTEN signaling or related pathways, such as PI3K-AKT, mTOR and NF-1 (e.g., rapamycin and everolimus), are potential candidates for treating several manifestations of autism(28).

The main goal of this study is to identify molecular mechanistic connections between the two groups of complex disorders. With this aim, we conducted meta-analyses of differential RNA expression of ASD brain tissues, and compared the dysregulated RNAs and related pathways with those involved in a collection of 22 tumor types and two non-cancer control diseases. Additionally, we employed the LINCS L1000 database(29, 30) to detect drugs with similar or opposite gene expression signatures to those of ASD and cancer(31). Finally, we specifically examined which elements of the PI3K-Akt-mTOR signaling axis (involving PTEN, FMR1, NF1, TSC1, and TSC2) were dysregulated jointly in ASD and cancer(32, 33).

## Materials and methods

### Data acquisition

Using the Gene Expression Omnibus (GEO, https://www.ncbi.nlm.nih.gov/geo/) and Array Express (AE, https://www.ebi.ac.uk/arrayexpress/) we retrieved RNA expression studies from ASD brain tissues and cancer. To apply uniform normalization methods to microarray raw data, we selected case-control datasets belonging to the most popular single channel array platforms from Agilent, Affymetrix, and Illumina. In the case of ASD, given the small number of studies available, an RNA-Seq dataset was also incorporated.

Three studies, which included ASD and control frontal cortex samples, were found and retrieved from public repositories or were obtained directly from the authors. The datasets of Chow(34) and Voineagu(35) (GSE28475 and GSE28521) were generated using the Illumina array platform HumanRef-8 v3.0, whereas Gupta’s dataset(36) was generated using Illumina’s HiSeq 2000 sequencing-technology (**Figure 1 A**). Cerebellar, temporal, and occipital cortex samples derived from the same set of patients were also available in GSE28521 and Gupta’s datasets. However, we focused on frontal cortex data to avoid introducing heterogeneity into the analysis due to tissue variability and because frontal cortex data was the represented by the highest number of samples. In the case of cancer datasets, we only used primary tumors and their healthy matched control tissues, and excluded other datasets and samples (i.e., metastasis and cell lines).

**Figure 1:**
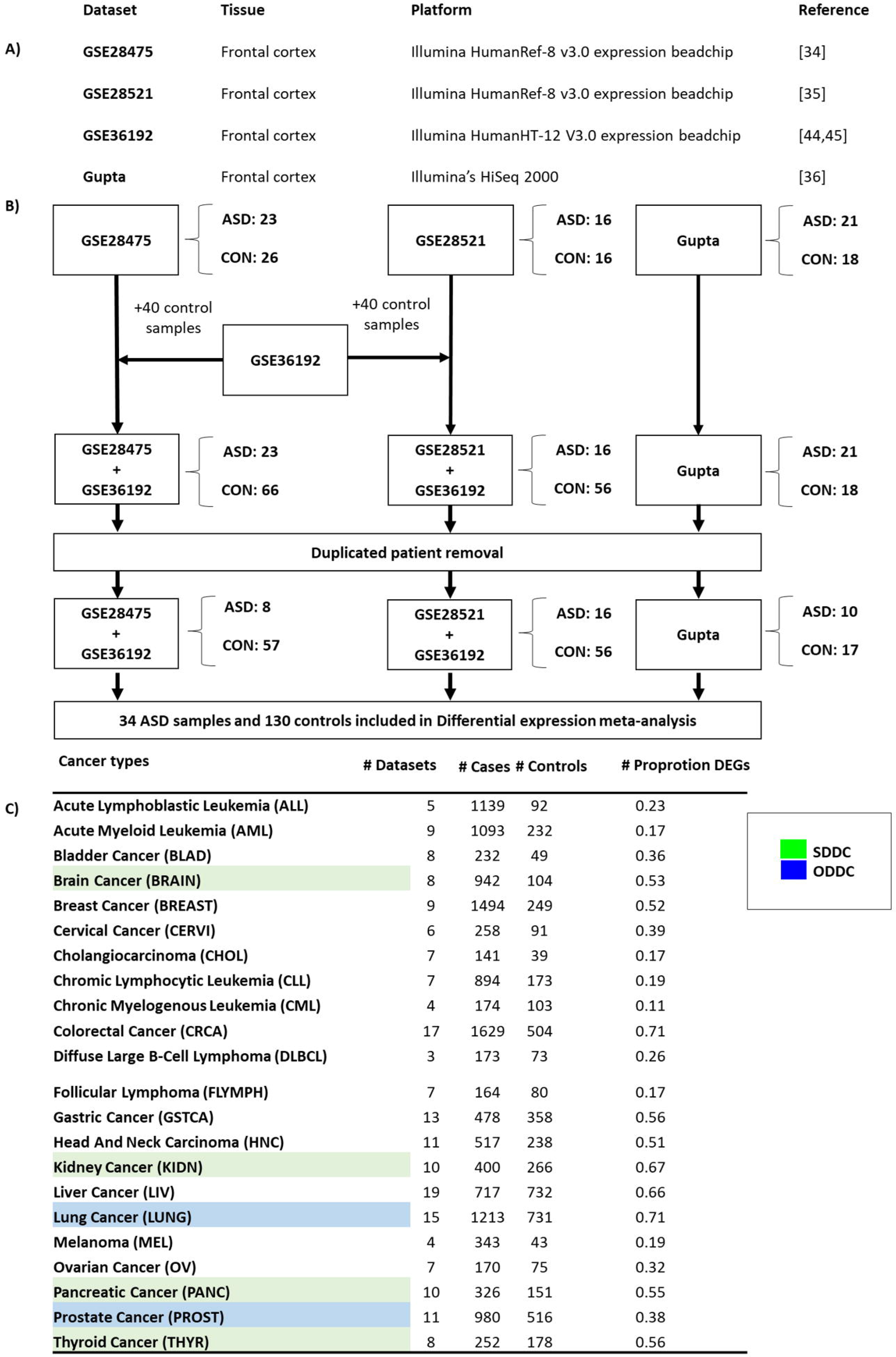
A) Table showing the datasets included in the ASD differential gene expression meta-analysis. B) Diagram depicting the workflow used to perform the ASD differential gene expression meta-analysis. C) Summary of the cancer types, number of datasets and samples included in each cancer-specific differential gene expression meta-analysis.

### Expression data preprocessing and normalization

Datasets generated using Affymetrix platforms were preprocessed as follows: CEL files were retrieved from GEO or AE, and the R packages oligo(37) and affy(38) were used to read the files and perform RMA normalization and summarization, which was followed by quantile between-sample normalization and log2 transformation. For Illumina platforms, non-normalized data were loaded to the R environment, and the Lumi(39) package was used to perform background correction using a normal exponential model fitting followed by quantile normalization and log2 transformation. Agilent data were preprocessed using the limma(40) package following the same preprocessing steps. In the case of RNA-Seq data, raw counts were loaded in the R environment. The Rlog function from the DESeq2(41) package was utilized to transform the RNA-Seq count distribution to a continuous distribution suitable for integration with the array data. In short, the Rlog function transforms count data into a continuous log2 scale distribution, minimizing the differences between samples for rows with small counts and normalizing the data with respect to library size. **Supplementary Figure 1** shows a comparison between two state of the art RNA-Seq specific differential expression methods and traditional limma analysis using Rlog transformed data suggesting that Rlog transformation renders the dataset suitable for inclusion with micro-array datasets.

To harmonize probe annotations between different dataset platforms, dataset-specific IDs were transformed into ENTREZ IDs using annotation packages. Probes targeting the same gene were collapsed using the collapseRows function from the WGCNA(42, 43) package selecting the MaxRowVariance method.

### Outlier exclusion

Potential outlier samples, defined using the following criteria, were removed from each dataset. We computed mean inter-array correlations prior to normalization for cases and controls independently. If the mean inter-array correlation within each group was lower than 0.9, we removed the sample showing the lowest mean inter-array correlations iteratively until a global correlation value of 0.9 was reached for both case and control groups. This method ensures that samples are not eliminated as outliers due to unbalanced case control designs while guaranteeing the elimination of samples with significant deviance from the group distributions. **Supplementary Table 1** shows the initial number of samples included in each study and the final number of samples after exclusion criteria were applied and outlier samples removed.

### Addition of control samples

The small number of samples found in the ASD brain transcriptomic studies limits the statistical power of differential expression meta-analysis. To enhance the power, searches were performed for additional datasets including frontal cortex control samples profiled with a compatible array platform. One study with such characteristics (GSE36192(44, 45) was found and retrieved. It included samples from the frontal lobe of the cerebral cortex profiled with Illumina HumanHT-12 V3.0. Then, we randomly included 80 frontal cortex control samples from GSE36192 to GSE28475 and GSE28475 (40 samples to each dataset) while maintaining balanced sex and age distributions between the cases and controls. No significant changes in sex, age of post mortem interval (PMI) distributions were present between cases and controls after control sample addition (**Supplementary Table 2**). For each dataset (GSE36192 and GSE28475) data from the original study and the 40 extra control samples were merged at the raw level. Then, a normal exponential background correction method was applied to the combined data followed by quantile normalization and log2 transformation using the lumi package(39). The combat function from the sva package(43) was finally applied to each preprocessed combination of one of the original datasets plus the control sample set in order to remove batch effects derived from different study origins (**Figure 1 B**).

### Removal of redundant patient samples

Currently published ASD brain transcriptomic datasets rely on a set of samples derived from a partially overlapping group of patients. Eleven ASD and one control sample derived from the same patients were included in both GSE28521 and Gupta’s dataset. Fourteen ASD and five control redundant samples were included in both GSE28475 and Gupta’s datasets. Ten ASD and four control redundant samples respectively were shared between GSE28475 and GSE28521. Nine ASD samples derived from the same patients were present in all three datasets. No common control samples were included in the three datasets (**Supplementary Figure 2**).

**Figure 2:**
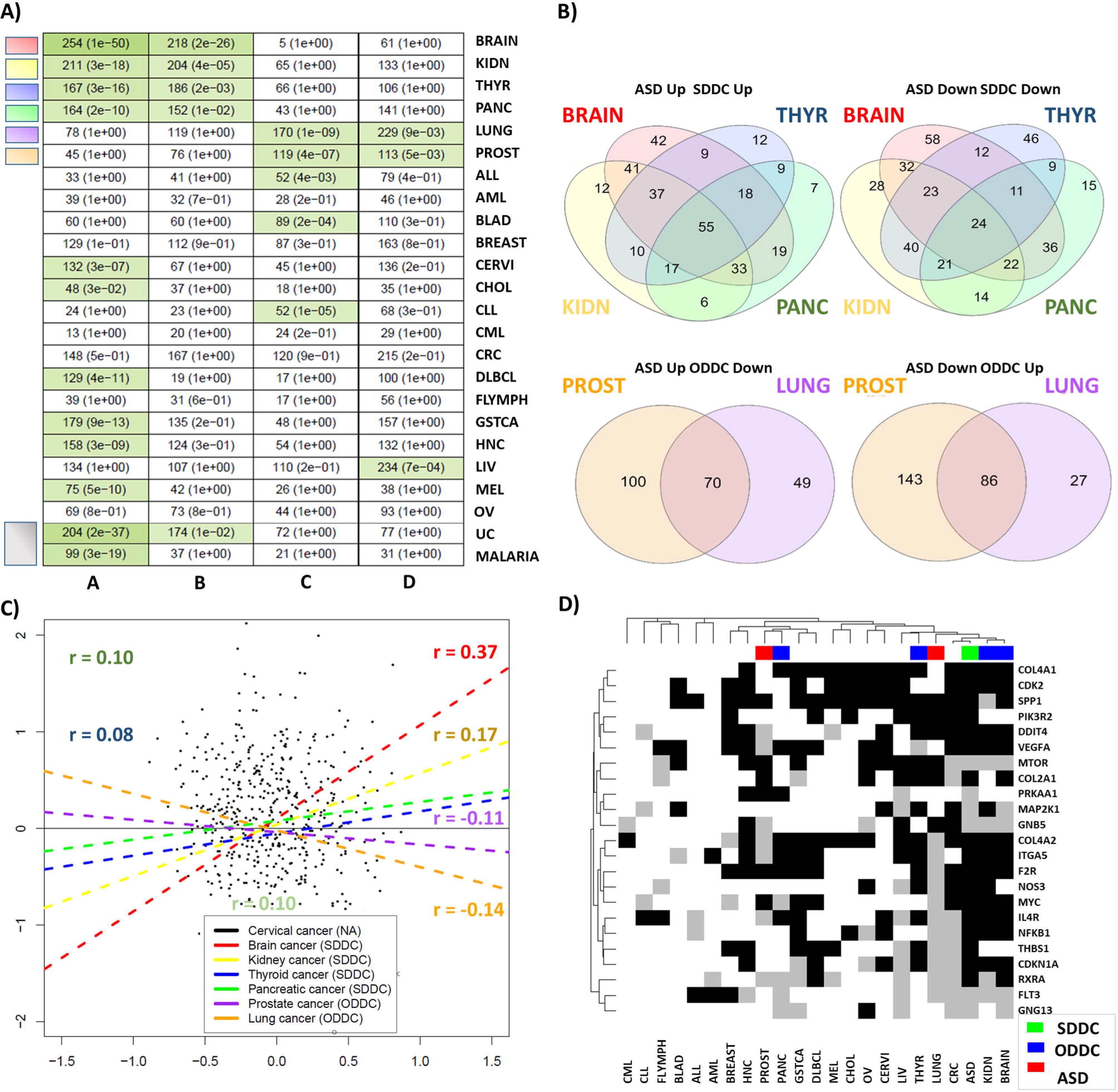
A) Table showing the significance of the intersections of upregulated and downregulated genes between ASD and the 22 cancer types included in our study, comprising, acute lymphoblastic leukemia (ALL), acute myeloid leukemia (AML), bladder cancer (BLAD), brain cancer (BRAIN), breast cancer (BREAST), cervical cancer (CERVI), cholangiocarcinoma (CHOL), chronic lymphocytic leukemia (CLL), chronic myeloid leukemia (CML), colorectal cancer (CRC), diffuse large b cell lymphoma (DLBCL), follicular lymphoma (FLYMPH), gastric cancer (GSTCA), head and neck carcinoma (HNC), kidney cancer (KIDN), liver cancer (LIV), lung cancer (LUNG), melanoma (MEL), ovarian cancer (OV), pancreatic cancer (PANC), prostate cancer (PROST), and thyroid cancer (THYR). Columns A, B, C, and D include the number of genes upregulated in both, downregulated in both, upregulated in ASD and downregulated in cancer, and downregulated in ASD and upregulated in cancer, respectively. Green cell colors indicate significant intersections (FDR corrected p-values from Fisher’s exact test lower than 0.05) with darker green tones indicating lower FDR corrected p-values. B) Venn diagrams showing the number of genes commonly deregulated in SDDCs and ODDCs. C) Scatter plots and correlation values, depicting the associations between ASD and all SDDC and ODDCs for cancer differential expression profiles. D) Heatmap showing the differential expression status of genes included in the KEGG hsa04151 pathways (PI3K-Akt signaling pathway), that were found to be differentially expressed in the ASD differential expression meta-analyses. White, gray and black cells indicate unaltered, downregulated and upregulated differential expression status, respectively.

Since patient redundancy could artificially inflate the number of differentially expressed genes yielded by differential gene expression meta-analysis, redundant samples were removed sequentially using the following criteria.

First, the R MetaQC package(46) was used to generate an index for the quality of each study. MetaQC integrates six quantitative quality control measures, appraising internal homogeneity of co-expression structure among studies, external consistency of co-expression patterns with a pathway database, and accuracy and consistency of differentially expressed gene detection or enriched pathway identification. For each dataset, the algorithm produces an index called standardized mean rank value (SMR) that can be interpreted as a relative measure of the quality of the study. SMR values were 1.17, 2.33 and 2.5 for GSE28521, Gupta, and GSE28475 respectively (**Supplementary Figures 3 A and 3 B**).

**Figure 3:**
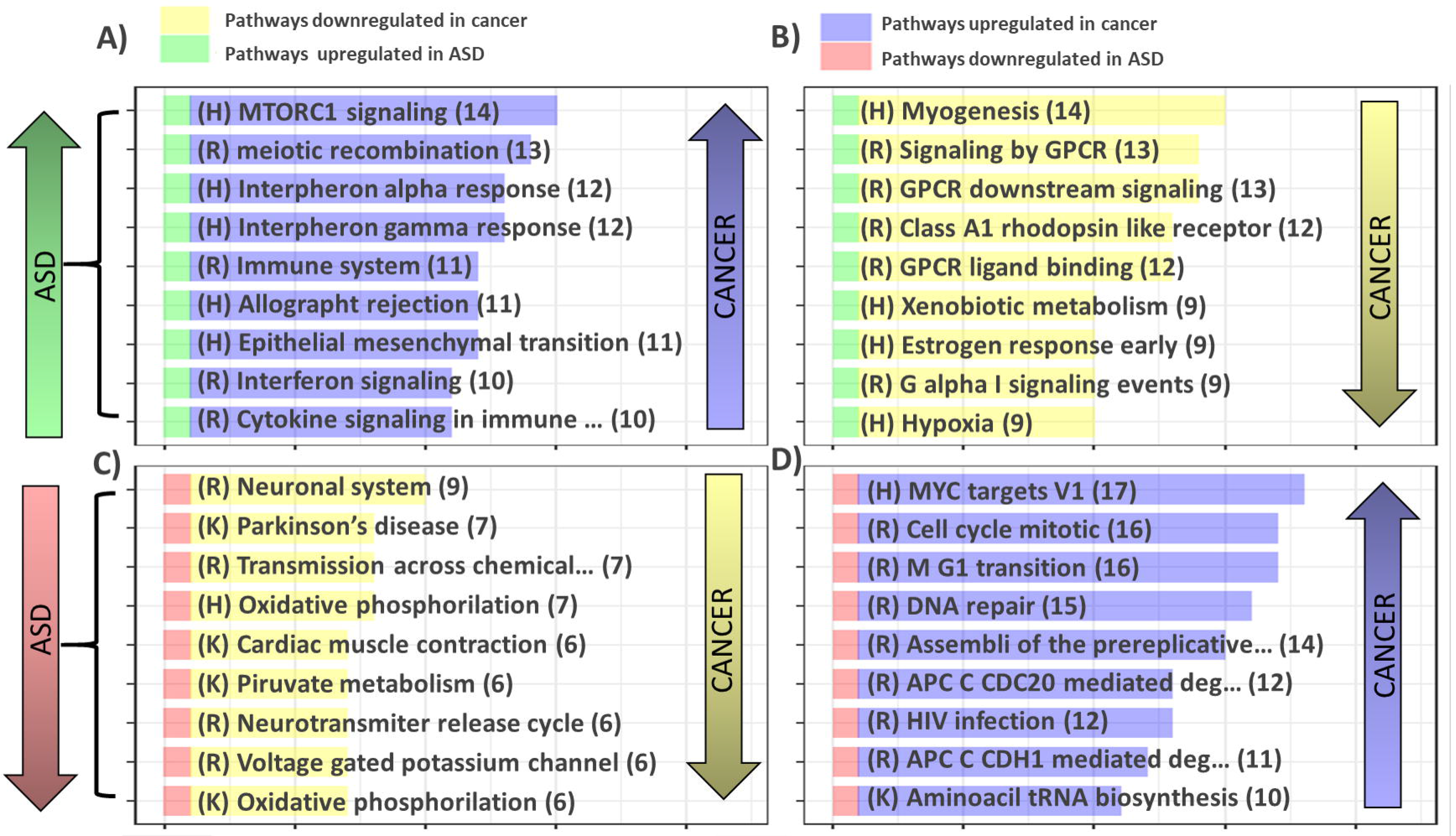
Top 10 ASD- and cancer-associated pathways extracted from 3 different molecular signature databases (Hallmarks, KEGG and Reactome). Yellow and purple segments indicate pathways downregulated and upregulated in cancer, respectively, whereas red and green segments denote pathways downregulated and upregulated in ASD, respectively. The length of the yellow and purple bars indicates the number of studied cancers that represent the reported direction of deregulation for this particular pathway.

Additionally, as an alternative quality metric, mean inter-sample correlations were computed for each dataset. GSE28521, Gupta, and GSE28475 showed mean inter-sample correlation values of 0.96, 0.95, and 0.91, respectively, which was in agreement with the quality ordering established by MetaQC. Using both criteria, we defined GSE28521 as the highest quality study, followed by Gupta, and GSE28475. To preserve the maximum number of samples in the highest quality studies, we kept all samples in the study that showed the lowest SMR value and the highest mean inter-sample correlation value (highest quality). Next, the samples derived from the same patients present in both the highest quality study and the study showing the next lowest SMR and the next highest mean inter-sample correlation (the second highest quality study) were removed from the second study. Finally, from the study showing the highest SMR value and the lowest mean inter-sample correlation (lowest quality), we removed the samples derived from individuals present either in the first or in the second highest quality studies (**Figure 1 B**).

After control sample addition and removal of duplicated individual samples, 34 non redundant ASD cases and 130 control samples distributed in the three datasets were available to perform differential gene expression meta-analysis. No significant differences in age, sex or PMI interval composition were found either between the cases and controls (p-value > 0.05) (**Figure 1 B**). **Supplementary Table 3** shows the samples included in the final ASD analysis and their associated covariates.

### Cancer and control diseases datasets

A total of 198 datasets from 22 different cancer types comprising 18736 samples, 13687 tumors and 5009 tissue-matched control samples were included in our cancer differential gene expression meta-analyses (**Figure 1 C**, **Supplementary Table 1**). The number of included datasets for each cancer type ranged from 3, in the case of diffuse large b-cell lymphoma, to 19 in the case of liver cancer. The sample sizes ranged from 180 in the case of cholangiocarcinoma to 2133 in the case of colorectal cancer. Malaria and ulcerative colitis were included as control diseases in order to evaluate the specificity of the associations between ASD and cancer. Ten ulcerative colitis datasets including 442 cases and 189 controls and three malaria datasets including 174 cases and 95 controls were used to perform differential gene expression meta-analyses.

### Differential gene expression meta-analyses

Differential gene expression meta-analyses are known to increase the statistical power and reduce the noise of gene expression measurements(47). For each disease, microarray meta-analyses were carried out independently using the approach developed by Choi et al(48) implemented in the MetaDE package(49). All meta-analyses were performed using random effect models, since moderate to high heterogeneity was expected given the biological and technical variability present in our data. The threshold of significance was set to a conventional level of 0.05. Thus, genes with a false discovery rate (FDR)-corrected p-value lower than 0.05 were considered differentially expressed.

### Comparison of differentially expressed gene profiles in ASD and cancer

The expression profiles of ASD and all studied cancer types were compared to evaluate the significance of the overlaps between differentially expressed genes, as previously described (50, 51). For each ASD-cancer pair, the significance of the four possible intersections formed by upregulated and downregulated genes was evaluated by means of one-tailed Fisher’s exact tests. The intersections were:

1. Genes upregulated in both autism and the selected cancer type (**Intersection A**),
2. Genes downregulated in both ASD and the selected cancer type (**Intersection B**),
3. Genes upregulated in ASD and downregulated in the selected cancer type (**Intersection C**), and
4. Genes downregulated in ASD and upregulated in the selected cancer type (**Intersection D**).

P-values were corrected by multiple testing using the FDR. Overlaps showing corrected p-values lower than 0.05 were considered significant. The background number of genes was set as the number of genes jointly studied in the two meta-analyses under consideration, which in turn depended on the platforms included in each meta-analysis. A cancer type was considered to be deregulated in the same direction as ASD when Intersections A and B were significant and Intersections C and D were not. These cancer types are referred to as same direction deregulated cancers (SDDCs) and could be candidates for direct comorbidity with ASD. Conversely, a cancer type was considered to be deregulated in the opposite direction from ASD when intersections C and D were significant but intersections A and B were not. These cancer types are referred to as opposite direction deregulated cancers (ODDCs) and could be candidates for inverse comorbidity with ASD.

An additional association analysis was performed on the differential expression profiles between all possible ASD and cancer pairs. Pearson’s correlation coefficients of the 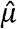 values obtained from each differential expression meta-analysis were computed. Positive correlations suggest similar patterns of differential expression while negative correlations would indicate opposite patterns.

### Gene set enrichment analysis

Gene set enrichment analysis was performed using Gene Set Enrichment Analysis (GSEA) (http://software.broadinstitute.org/gsea/msigdb) in order to detect functional categories enriched in upregulated or downregulated genes. Z-values produced as output in each differential expression meta-analysis were employed as an ordering factor. For each disease, enrichment calculations were carried out using different molecular signature databases, namely, Hallmarks (H), Canonical pathways (C2), and Gene Ontology (GO). A significance threshold of 0.05 was defined for the corrected p-value generated by the GSEA algorithm when selecting enriched functional categories.

For enrichment of gene sets placed on the intersections, a traditional overrepresentation analysis was performed using g:Profiler, an online tool for functional profiling of gene lists from large-scale experiments, through the interface R package gProfileR(52).

### LINCS-based analysis and drug set enrichment analysis

LINCS L1000(27, 29, 53) (http://amp.pharm.mssm.edu/L1000CDS2/#/index) comprises a collection of 230,556 gene expression profiles of cancer cell lines perturbed by small molecules and genetic constructs. Here, a subset of 29,157 small molecule perturbations that was included in a custom drug classification partially based on the anatomical therapeutic chemical classification system (ATC) was selected and employed to perform drug set enrichment analyses for each studied condition as previously described in Sanchez et al(51).

A list of genes ranked based on z-values derived from the differential gene expression analysis was generated for each disease. Then, each ranked gene list was used to compute cosine distances with each of the 29 157 perturbations included in our drug classification using the R ccmap package(54). This method produces a list of perturbations or drugs ordered by its cosine distance with the target disease. Positive cosine distances indicate that a particular small molecule or drug produces a differential expression profile that mimics or resembles the differential expression profile of the disease under consideration, whereas negative cosine distances suggest that a particular small molecule or drug produces a differential expression profile that reverses the target disease profile.

Finally, the list of small molecules or drugs ordered by their cosine distance with the differential expression profile of a particular disease, was used to detect enrichment in drug sets using a GSEA-based enrichment method implemented in the fgsea package(55). The algorithm reveals whether a particular drug set is preferentially located at one of the extremes of the ranked list of drugs associated with each disease. Significant placement of a particular drug set at the top of the distribution suggests that it produces an effect that mimics the transcriptomic changes found in the disease under consideration. Conversely, significant placement of a particular drug set at the bottom of the perturbation distribution suggests that it produces an effect that reverses the transcriptomic changes found in the disease under consideration. A conventional FDR value of 0.05 was selected as a threshold

We carried out this analysis for all ASD and cancer differential expression profiles. Finally, the results were compared between ASD and each tumor type.

## Results

### ASD differential gene-expression meta-analysis and gene set enrichment analysis (GSEA)

A total of 13,699 genes were tested for differential expression, yielding 1,055 differentially expressed genes (DEGs) in ASD patients relative to controls below an FDR threshold of 0.05. Of these DEGs, 450 were upregulated and 605 were downregulated (**Supplementary File 1**).

Gene set enrichment analysis (GSEA) suggested that genes upregulated in ASD are mainly associated with immune system-related processes, including cytokine production, inflammatory response, leukocyte activation, NFKB signaling, interferon response and complement reaction. Cell death regulation, cell adhesion, P53 signaling, and extracellular matrix organization were also enriched in upregulated genes. Genes downregulated in ASD samples were mainly associated with oxidative phosphorylation, ATP metabolism and lactic acidosis. Neuronal system functions, such as GABA synthesis, reuptake, and degradation plus proteasome pathway related processes, were also enriched in genes downregulated in ASD samples compared to controls (**Supplementary File 2**).

### Cancer data analysis

We found a very high proportion of differentially expressed genes in our cancer meta-analysis results, with values ranging from 11% to 71% of the total number of tested genes in chronic myeloid leukemia (CML) and colorectal cancer, respectively (**Figure 1 C**, **Supplementary File 4**). These proportions are compatible with previous findings for differential gene expression analysis of TCGA RNA-Seq cancer data(56), where the percentage of differentially expressed genes ranged from 32% in the case of bladder cancer to 72% in the case of breast cancer. The number of differentially expressed genes found in our analysis was correlated to the number of included studies (r = 0.76) and samples (r = 0.53), and the minimum weighted mean difference 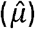 for a gene detected as significantly differentially expressed in a particular meta-analysis negatively correlated with the number of included studies (r = −0.73) and samples (r = −0.67). This finding indicates that as more studies were introduced in the meta-analyses, genes with smaller but consistent differences in expression were detected as significantly deregulated.

Enrichment analysis showed that pathways associated with cell cycle such as, mitotic phase transition, DNA synthesis and repair, and telomere extension, were commonly upregulated in most of the cancer types (68%). Interestingly, leukemias and lymphomas did not show changes in mitosis related pathways (**Supplementary Figures 4 and 5**). The most common downregulated pathways among cancers were related to calcium, G-protein-coupled receptors (GPCR) signaling, and fatty acid metabolism which were downregulated in between 40 and 50% of the studied cancers.

### Autism and cancer expression deregulation profile comparisons

To investigate whether the transcriptomic deregulations observed in particular cancer types showed direct or inverse patterns of association with ASD, all possible ASD and cancer pairs were subjected to intersection and correlation analysis (**See Methods**). Four cancer types (brain, kidney, pancreatic, and thyroid cancer) presented differential gene expression profiles that were significantly deregulated in the same direction as ASD below an FDR corrected p-value threshold of 0.05. These tumor types are referred to as same direction deregulated cancers (SDDC). Two tumor types (lung and prostate cancer) showed differential expression profiles deregulated in the opposite direction from frontal cortex samples of ASD patients. These types are referred to as opposite direction deregulated cancers (ODDC). No association was present between ASD and the rest of the cancers. (**Figure 2 A**).

Two hundred and fifty-four genes were found to be jointly upregulated in ASD and brain cancer and were enriched in immune system and cell death related processes. Two hundred and eighteen genes were found to be jointly downregulated in ASD and brain cancer. Enrichment in neuron and synapse related genes was found in this set of genes. Similar enrichment results were observed in the analysis of the 164 and 152 genes jointly up- and downregulated in ASD and pancreatic cancer. Kidney cancer and ASD presented 211 and 204 genes jointly up- and downregulated respectively. Shared upregulated genes between ASD and kidney cancer were also heavily enriched in immune related processes and cell death. Jointly downregulated genes in ASD and kidney cancer were enriched in mitochondrial functions and ATP synthesis. Similar results were obtained in the analysis of the 167 and 186 genes jointly up- and downregulated, respectively, in ASD and thyroid cancer, showing strong enrichment in immune system and mitochondrial function related genes in the joint upregulated and downregulated gene sets respectively.

One hundred and seventy genes were found to be jointly upregulated in ASD and downregulated in lung cancer and were enriched in immune system processes and cell death among others, whereas 229 genes were found to be downregulated in ASD and upregulated in lung cancer showing enrichment in functions related to mitochondrial function. One hundred and nineteen were found to be upregulated in ASD and downregulated in prostate cancer which were also enriched in focal adhesion, cell death and immune system processes whereas 113 gees were found to be downregulated in ASD and upregulated in prostate cancer which were enriched in mitochondrial related functions.

**Supplementary Table 4** and **Supplementary File 4** show the genes placed in the described intersections and the results of the overrepresentation-based functional analysis.

To determine the degree of homogeneity within SDDC and ODDC groups, we compared the content of the previously described intersections in each group. A total of 55 (17%) and 24 (6%) genes were jointly up- and downregulated respectively in the four SDDCs and ASD. Seventy (31%) genes were upregulated in ASD and downregulated in both ODDCs, whereas 86 (35%) were downregulated in ASD and upregulated in both ODDCs (**Supplementary Table 5**, **Figure 2 B**).

To evaluate the level of specificity of the reported associations between ASD and cancer and determine if associations with previous epidemiological confirmation translate in same or opposite direction deregulation patters, ASD and cancer differential expression profiles were compared to two control diseases, ulcerative colitis (UC) and malaria. Ulcerative colitis has been shown to have direct comorbid associations with both ASD and colorectal cancer (CRC) (57, 58). UC differential expression profile was found to be deregulated in the same direction as both, ASD and colorectal cancer (**Supplementary Figure 7 A**). Nine other cancer types were found to be deregulated in the same direction than UC including all SDDCs. Prostate cancer, ALL, and CLL were found to be deregulated in opposite directions as UC. No reports investigating associations between malaria and ASD or cancer have been published to the date. Our results showed no transcriptomic associations between malaria and ASD differential expression profiles. Thyroid cancer was found to be deregulated in the same direction as malaria whereas ALL and CLL were found to be deregulated in opposite directions (**Supplementary Figure 7 B**).

Complementarily, we computed Pearson’s correlation between the differential expression profiles of each possible pair of ASD and cancer to quantify the degree of association between them. SDDCs showed positive correlations with ASD. Brain cancer was the cancer type that showed the highest correlation values (r = 0.37, p < 0.05), and it was followed by kidney, thyroid and pancreatic cancer (r = 0.17, r = 0.10, and r = 0.08, respectively, FDR < 0.05). ODDCs differential gene expression profiles presented negative correlations with ASD. Lung and prostate cancer showed significant negative correlations (r = −0.14 and r = −0.11, respectively, FDR < 0.05). All other cancer types presented correlation absolute values lower than 0.1 (**Figure 2 C**). Associations showing the lowest FDR corrected p-values in the intersection analysis tended to present the strongest Pearson’s correlations.

Partition Around Medoids (PAM) cluster analysis was carried out on the differential expression profiles of ASD and the 22 tumor types. Silhouette analysis was first applied to determine the optimum number of clusters. The five groups partition showed the highest average silhouette value suggesting that 5 was the optimum number of clusters. However, the average silhouette value was low in all cases indicating the absence of substantial structure. Results for different number of partitions can be found in **Supplementary Figure 5**. Overall, ASD tends to cluster together with brain cancer. Cancers include in the ODDC and SDDC groups tended to group together in the same cluster, indicating that their differential expression profiles were more similar between them compared to other cancer types.

A theoretical overall cancer gene expression profile was constructed by averaging the differential expression profiles of all studied cancer types. No association (r = 0.05) was observed between ASD and this theoretical overall cancer profiles.

### PI3K associated genes

Given the pivotal role that PI3K/AKT/MTOR plays in both ASD and cancer, we studied the differential expression status of the genes included in KEGG’s hsa04151 pathway (PI3K-Akt signaling pathway). Twenty-five genes out of 272 genes belonging to hsa04151 (Fisher’s exact test p-value = 0.21) were found to be deregulated in ASD below an FDR threshold of 0.05, including the core pathway gene MTOR, which was found to be downregulated in ASD. Twenty-three out of 25 genes were present in the meta-analysis of ASD and the 22 cancer types.

To determine the degree of similarity among the deregulation patterns of the 23 PI3K-associated genes observed in ASD and present in all meta-analysis cancer results, we performed hierarchical clustering using different distance measures using as an input a matrix containing discrete values for each gene representing upregulation, downregulation and normal expression status (**Figure 2 D**). Brain and kidney cancer clustered together with ASD with all distance measures employed suggesting common patterns of changes in this subset of genes belonging to the PI3K-Akt signaling pathway. (**Supplementary Figure 6**). In particular, F2R, MYC, NFKB1, VEGFA, DDIT4, CDKN1A, CDK2, ITGA5, COL4A1, COL4A2, and IL4R were upregulated in ASD and brain and kidney cancer, while MTOR, FLT3, and GNB5 were found to be downregulated in these three diseases.

### Pathway enrichment analysis comparisons and LINCs drug set analysis results

To sketch the landscape of global common biological pathway dysregulation between ASD and cancer, we carried out functional analysis of the differential expression meta-analyses results for each included disease. To this end, GSEA and LINCS drug set enrichment analysis were performed as described in Methods. Immune system associated pathways, such as interferon alpha and gamma signaling, IL6 JAK STAT3 signaling, TNFA signaling through NFKB and MTORC1 signaling, were found to be upregulated in both ASD and 55%, 55%, 41%, 41% and 63% of cancer types, respectively. Jointly downregulated pathways between ASD and cancer were mainly associated with neuronal system genes, oxidative phosphorylation and ATP synthesis in 41% and 31% of cancers respectively. However, oxidative phosphorylation was also found to be upregulated in a subset off cancer types indicating differences in the energy metabolism abnormalities found in different tumor types. Processes downregulated in ASD and upregulated in cancer included MYC targets, DNA repair, HIV infection and proteasome activity in 77%, 68%, 55%, and 46% of the studied cancers, respectively, whereas GPCR signaling and myogenesis are examples of pathways that were upregulated in ASD and downregulated in 59% and 63% of cancers, respectively (**Figure 3**).

Drug sets commonly linked to ASD and cancer were also examined. The results suggest that treatment with mTOR inhibitors, such as everolimus, sirolimus, and temsirolimus, produce differential expression profiles that mimic the differential expression profile found in ASD while reversing the differential expression profiles found in most cancer types, excluding brain, kidney, thyroid, and pancreatic cancer, the four SDDCs.

STAT signaling inhibition by niclosamide produces differential expression profiles that mimic the ASD DEG signature while reversing the differential expression profiles of 40% of the studied cancers. Proteasome inhibitors and histone deacetylase inhibitors, such as bortozemib, entinostat and vorinostat, also mimicked ASD differential expression profile while reversing the differential expression profiles of 40% of the studied cancers (**Figure 4**).

**Figure 4).**
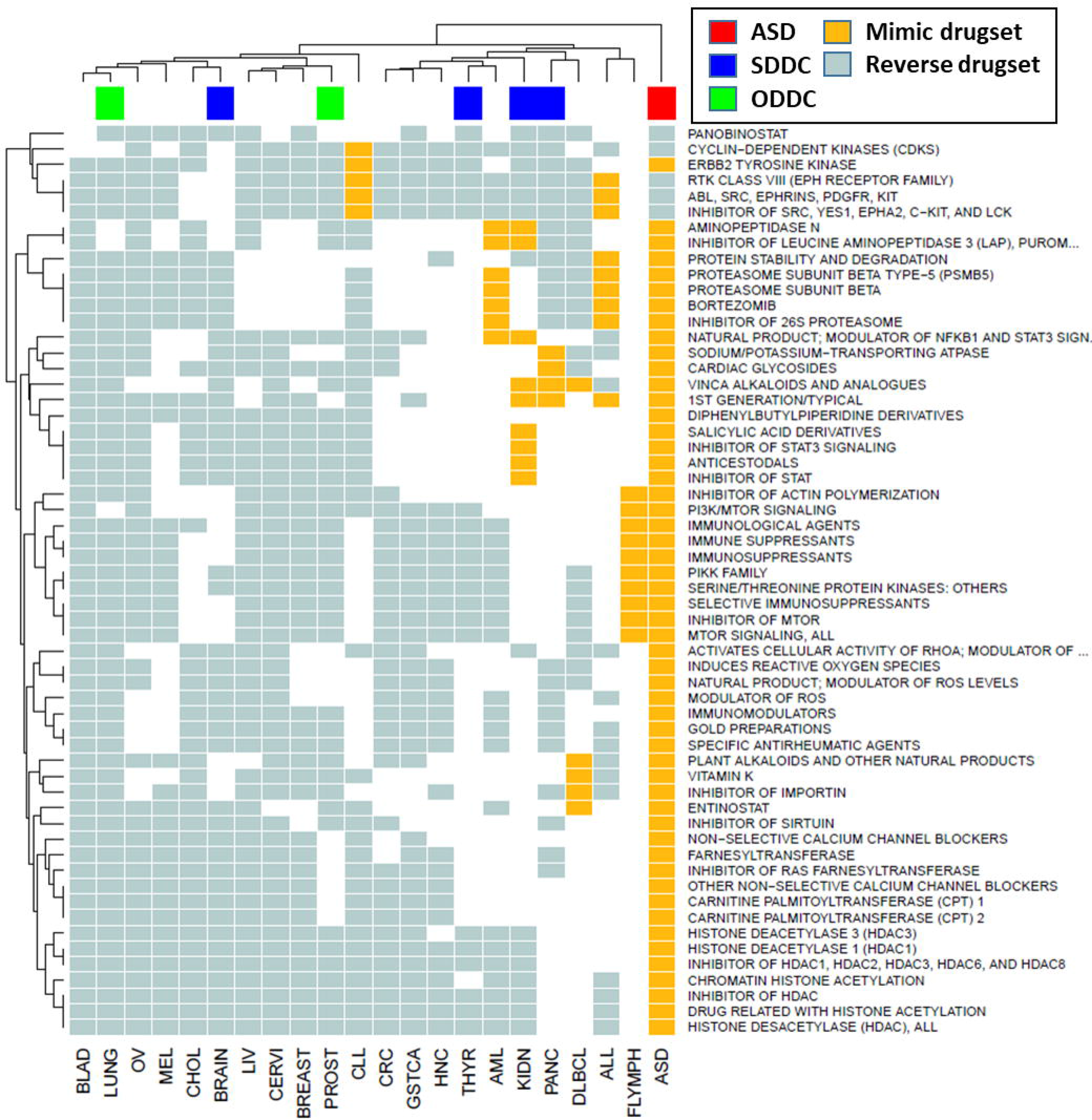
LINCS L1000-derived top related drug sets. Gold cells represent drug sets that produce differential expression profiles that mimic the differential expression profiles found in the disease of interest, light blue cells indicate drug sets that generate differential expression profiles opposite to those found in our disease of interest. Green, blue and red bars located on top of the heat map indicate ODDCs, SDDCs and ASD membership, respectively.

Restricting the analysis to cancers significantly associated with ASD, we observed that pathways jointly affected in ASD and SDDCs were mainly dysregulated in the same direction, i.e., they were upregulated or downregulated in both diseases. Their proportions in brain, kidney, thyroid, and pancreatic cancer were 85%, 89%, 88% and 65%, respectively, while pathways jointly affected in ODDCs and ASD were mainly deregulated in opposite directions (96% and 95% for lung and prostate cancer, respectively) (**Supplementary Figure 8**). Upregulated pathways shared by SDDCs and ASD were fundamentally linked to immune system-related processes. Shared downregulated pathways were implicated in oxidative phosphorylation, GPCR signaling and neuronal system genes. ODDCs upregulated pathways included cell cycle and DNA repair pathways. Contrary to what we observed in SDDCs, oxidative phosphorylation-related pathways were also upregulated in both lung and prostate cancer, indicating heterogeneity in energy metabolism abnormalities in different cancer types. Apoptotic, focal adhesion, and MAPK pathways were downregulated in ODDCs and upregulated in ASD. Finally, the MTORC1 pathway was found to be deregulated in ASD, SDDCs and ODDCs (**Figure 5**).

**Figure 5).**
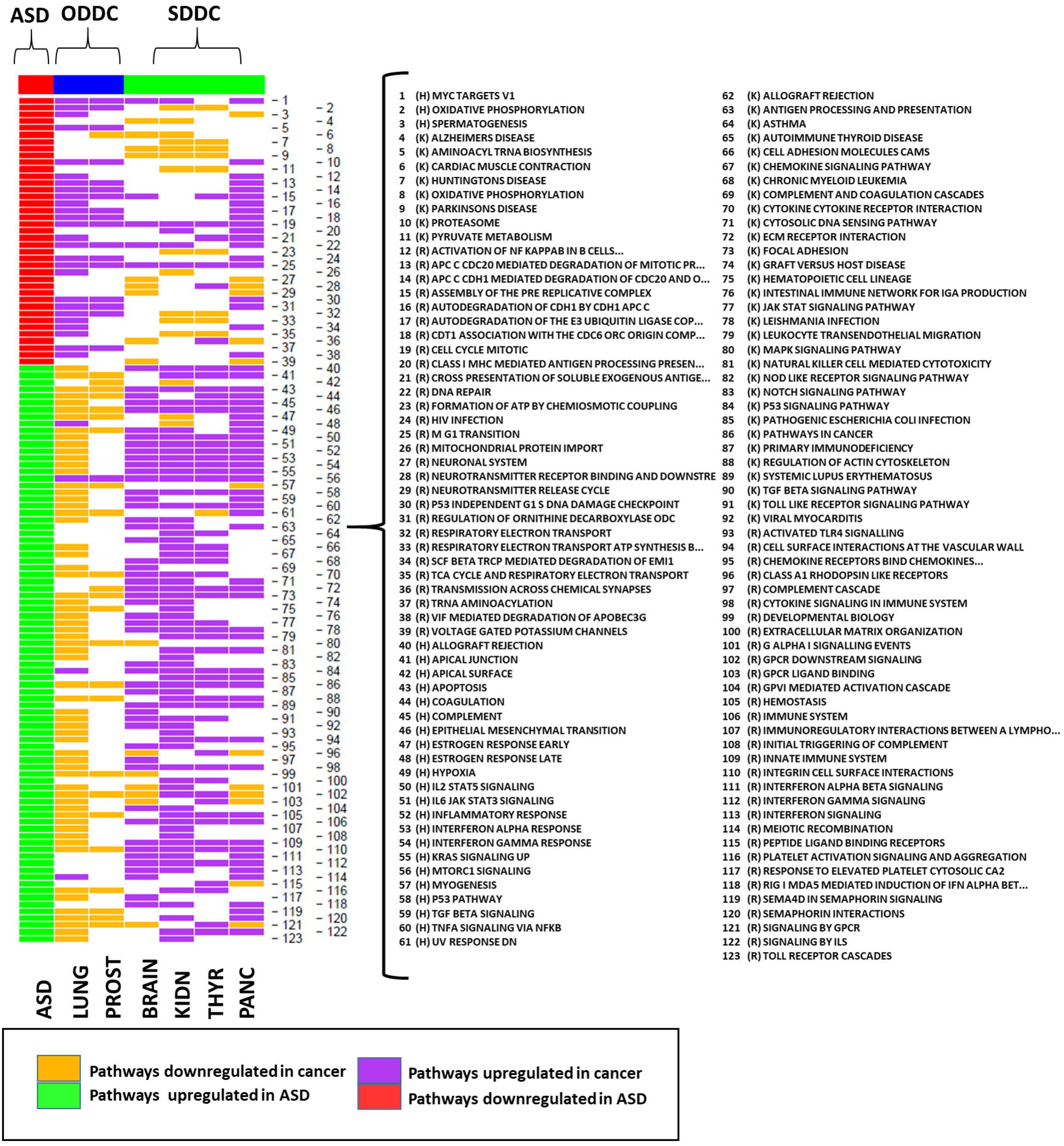
Heatmap showing the pathways altered in ASD, SDDCs and ODDCs. Yellow and purple cells indicate pathways downregulated and upregulated in cancer, respectively, red and green cells denote pathways downregulated and upregulated in ASD, respectively.

## Discussion

This is the first study aiming to explore the molecular associations between ASD and cancer at a transcriptomic level. We found positive patterns of association between ASD and four cancer types (brain, kidney, thyroid, and pancreatic) and negative patterns of association between ASD and two cancer types (lung and prostate). Brain cancer and kidney cancer showed the strongest transcriptomic associations with ASD in both intersection and correlation analyses. This observation is in agreement with previous epidemiological data reporting an increased risk of both benign and malignant brain neoplasms in patients with ASD(10). Interestingly, the same work also noted an increased risk of congenital malformations of the urinary system in autistic individuals, including medullary sponge kidney and the presence of accessory kidneys. Epidemiological associations between urogenital system tumors and ASD have also been reported(11).

The ASD differential expression results included genes that have previously been linked to both ASD and cancer(12, 24). For example, CUL3, a component of the multiple cullin ring ubiquitin-protein ligase complex(24), was downregulated in our ASD analysis. Furthermore, nine oncogenes present on the gene list compiled by Darbro(12) were found to be deregulated in ASD. Seven were found to be upregulated (ABL1, MYC, NFKB2, PIM1, PPARG, and BCL6) and 2 downregulated whereas two were found to be downregulated (FLT3 and MAP2K1).

On the one hand, a number of pathways were found to be commonly deregulated in different directions in ASD and several cancer types. Histone deacetylase activity, GPCR signaling, proteasome function, MYC targets, and cell cycle processes are representative examples. Some of the enumerated biological functions have previously been related to both ASD and cancer(59–64). These abnormalities could help explain putative inverse comorbid associations between ASD and cancer.

On the other hand, some biological processes were found to be deregulated in the same direction in both ASD and SDDCs, providing theoretical support for hypothetical direct comorbid associations between ASD and cancer. For instance, in agreement with previous data(65–70), our analysis suggests the presence of brain inflammation in ASD patients. Inflammatory processes are well-established drivers of carcinogenesis(71, 72) and are a factor that exerts direct influence on cancer-related features, such as proliferation, survival, and migration(72). In further support of this hypothesis, indicators of ongoing inflammation were observed in several cancers, including all tumors classified as SDDCs.

Different degrees of mitochondrial activity impairment were also observed as a shared trait between ASD and SDDCs. These changes were more evident in kidney and thyroid cancers, where oxidative phosphorylation, mitochondrial electron transport chain and ATP synthesis-related genes, including ATP50, ATP5F1, OGDHL, ATP5J, CYC1, PFKM, UQCRFS1, NDUFB6, NDUFB2, NDUFAF1, NDUFV1, DLD, and COX7B, were found to be jointly downregulated with ASD. Oxidative phosphorylation impairment, mitochondrial dysfunction and increased oxidative stress are distinctive features of autistic brains(73, 74). Some studies have suggested that genes regulating these processes are highly enriched in parvalbumin GABAergic interneurons of the forebrain, a cell type that has been implicated in multiple murine ASD models and in humans with ASD(75). Higher rates of glycolysis and suppression of mitochondrial function are traits commonly observed in cancer cells. Although advances in the understanding of cancer metabolism depict oxidative phosphorylation impairment as a more complex phenomenon than previously thought(76), our data suggest that this function is commonly impaired in at least a subset of tumor types. However, some cancer types, such as lung cancer showed opposite patterns of deregulation of mitochondria and ATP synthesis related genes, highlighting the heterogeneity present in cancer energy metabolism. In addition, there is evidence indicating that inflammation and oxidative phosphorylation may have a synergic effect. Cytokines, and particularly, tumor necrosis factor alpha (TNFα), impair mitochondrial oxidative phosphorylation and ATP production and increase reactive oxygen species (ROS), which in turn can increase mitochondrial injury and trigger mitochondrial content release to the cytosol, amplifying the inflammatory process(77). This interplay between the two processes may increase the risk of tumor development.

The PI3K/AKT/MTOR axis is an important target for molecular abnormalities in both ASD and cancer, which makes it a good candidate to modulate putative comorbid ASD and cancer associations. Our results showed that ASD patients presented patterns of dysregulation in this axis that are more similar to those observed in brain and kidney cancer than to any other tested studied cancer. Furthermore, GSEA and LINCS analyses suggested that the pathway is affected in ASD and a subset of cancers. However, given its complex nature(78, 79), which includes crosstalk with other signaling pathways and the presence of feedback loops, it is difficult to state whether the observed results are indicators of pathway activation or inhibition. Interestingly, ASD idiopathic cases and monogenic diseases related to autism have been linked to both higher and lower activity of the PI3K/AKT/MTOR axis(80–82). The PI3K/AKT/MTOR axis is one the most frequently altered pathways in human tumors and directly participates in the regulation of many cancer hallmarks(83). Moreover, it regulates several key events related to both inflammatory response, oxidative phosphorylation, and mitochondrial function(84–87). Our observations are in agreement with a recent review highlighting the importance of the PI3K/AKT/MTOR axis and mitochondrial abnormalities as potential modulators of ASD and cancer associations(23).

The analysis regarding the two control diseases, UC and malaria, showed that previously reported direct epidemiological associations between diseases translate into similar patterns of transcriptomic deregulation between them. The increased risk of UC in ASD patients observed at a population level(57) was followed by significant same direction deregulation patterns between both diseases at a transcriptomic level. Similarly, significant same direction transcriptomic changes were observed between UC and colorectal cancer (CRC), two diseases with a known direct epidemiologic link. Furthermore, intersection analysis of UC and cancer revealed that UC was positively associated with multiple cancer types including all SDDCs and inversely associated with three cancer types including prostate cancer (ODDC). These observations indicate that the transcriptomic associations between ASD and cancer suggested by our analysis are not ASD-specific and could be shared by other diseases showing similar gene expression deregulation patterns.

Finally, we recognize some important limitations. First, the number of available datasets with gene expression data derived from autistic patient brains is scarce as it is the number of included samples in each dataset. This fact has two main consequences. On one hand, it undermined the statistical power of differential expression meta-analysis. Furthermore, it impedes patient stratification, which would be advisable given the intrinsic heterogeneity of ASD. It is reasonable to expect that different subgroups of ASD individuals present different patterns of association with cancer. We are aware that several studies, including blood-derived transcriptomic profiles of autistic patients, have been published to date; however, it still is an open question as to whether differential expression profiles derived from peripheral tissues can be used as a proxy to detect molecular abnormalities directly linked to disease physiopathology. Second, the analysis of transcriptomes is often not enough to detect whether particular biological processes were activated or inactivated, this imposes a limit to the conclusions that can be drawn from our results. Finally, although scientific evidence is starting to accumulate in favor of the presence of comorbid associations between ASD and cancer, more population and molecular studies are needed to confirm or refute competing hypotheses.

In summary, immune-related processes, mitochondrial dysfunction, and PI3K/AKT/MTOR signaling are biological processes that have been independently associated with both ASD and cancer. We have described the presence of a variable degree of changes in these pathways in SDDC, including brain and kidney cancer, the two cancer types showing the strongest associations with ASD in our intersection analysis. This observation makes these pathways good candidates to intervene to modulate putative direct comorbid associations between them. In addition, we report opposite direction associations between ASD and particular cancer types and pathways that may represent underlying molecular substrates of theoretical inverse association between ASD and cancer. These findings show a complex interplay between potential comorbid associations of ASD and cancer, and highlight the importance of further research at epidemiological, genetic, and molecular level.

## Acknowledgements

Professor Rafael Tabarés-Seisdedos and Jaume Forés were supported in part by grant number PROMETEOII/2015/021 from Generalitat Valenciana and the national grant PI17/00719 from ISCIII-FEDER.

